# Discovery of BilV reveals a multienzymatic basis for bilirubin reduction across vertebrate gut microbiomes

**DOI:** 10.64898/2026.06.01.729425

**Authors:** Angela K. Jiang, Maggie R. Grant, Gabriela Arp, Keith Dufault-Thompson, Alexandra M. Clarke, Yue Li, Danielle Lehman, Alan K. Jarmusch, Brantley Hall, Xiaofang Jiang

## Abstract

Gut bacteria reduce bilirubin to urobilinogen, allowing it to be excreted through feces and urine, but studies have long noted a heterogeneous mixture of partially reduced bilirubin-derived intermediates, suggesting that multiple enzymes are involved. Here we identify bilirubin vinyl reductase (BilV), a novel Old Yellow Enzyme family reductase encoded in the genomic neighborhood of the known bilirubin reductase (*bilR*). Using heterologous expression and LC-MS/MS, we show that BilR acts on the methine bridges in the bilirubin reduction pathway; co-expression with BilV enables vinyl-group reduction and complete conversion to urobilinogen. In bacterial genomes, *bilV* co-occurs primarily with the *bilR*-insertion subtype and is largely absent alongside *bilR*-short. Analysis of 1,197 gut metagenomes across 14 vertebrate species reveals that this differential co-occurrence shapes pathway availability across hosts: carnivores and omnivores carry balanced *bilR* and *bilV*, whereas avian microbiomes, dominated by *bilR*-short, are depleted for *bilV*. These findings establish that bilirubin reduction to urobilinogen involves two enzymes with complementary regioselectivity, and that their distribution across vertebrate gut microbiomes varies in concert with host bile pigment chemistry.

## Introduction

As part of the natural turnover of hemoglobin and hemoproteins, heme is produced, incorporated, and then broken down into excretable waste products. In mammals, heme oxygenase converts heme to biliverdin and biliverdin reductase converts biliverdin to bilirubin IXα, which is conjugated in the liver and excreted via bile into the gut1. The final clearance steps are performed by anaerobic intestinal bacteria, which reduce bilirubin to urobilinogen and stercobilinogen for excretion in urine and feces, respectively2. When this microbial process is disrupted in neonates3 and germ-free animals1,4, bilirubin may re-enter enterohepatic circulation leading to hyperbilirubinemia, which in extreme cases, may lead to neurological damage5,6. Despite the clinical significance of the heme degradation process, the bacteria and genes involved in the bilirubin reduction pathway have only recently started to be identified3.

The conventional microbial pathway assumes a linear sequence of reductions proceeding from bilirubin to mesobilirubin to urobilinogen and finally stercobilinogen4. However, biochemical studies of fecal bacterial cultures indicate that this process is variable. Mass spectrometry revealed that when bilirubin served as substrate, a less-saturated, presumably monovinyl species (MW: 588) appeared alongside the fully reduced diethyl form (MW: 590), and dihydromesobilirubin (methylene-mesbolirubin) was hypothesized to be a key intermediate in the pathway7. These observations were structurally confirmed by NMR and corroborated by chromic acid degradation of urobilinoids produced from bilirubin in fecal bacterial cultures8,9. In addition, a range of partially reduced intermediates have been detected in bacterial cultures and human urine4,9. These studies indicate that the bacterial reduction of bilirubin and its derivatives might not be a fixed sequential process, leading to a mixture of end products. As such, it is possible that bilirubin reduction is a multienzymatic process, rather than the assumed single-enzyme reduction.

The bilirubin reductase enzyme, BilR, an FMN-dependent Old Yellow Enzyme (OYE) family oxidoreductase, was the first enzyme in the bilirubin reduction pathway to be characterized3. BilR reduces bilirubin and was found to be predominantly encoded by gut-associated Firmicutes species3. An analysis of the evolution of BilR demonstrated that the family contains three distinct clades (BilR-basal, BilR-short, and BilR-insertion) with differences in domain architecture10. Further analysis of over 1,000 vertebrate gut microbiomes revealed that BilR is enriched in the large intestines of mammals and chickens, and that the BilR clades have distinct distribution patterns associated with host-diet and gastrointestinal anatomy10. As birds have low hepatic biliverdin reductase activity, they primarily excrete biliverdin rather than bilirubin into their intestines11, making the presence of bilR genes in avian gut microbiomes unexpected. Either BilR acts directly on biliverdin as an alternative substrate, or an uncharacterized enzyme first converts biliverdin to bilirubin in the avian gut. Either scenario points to enzymatic chemistry not previously recognized in the bilirubin reduction pathway.

In this study, we present the identification and characterization of a novel Old Yellow Enzyme family reductase, BilV (bilirubin vinyl reductase), which reduces vinyl groups within bilirubin-derived intermediates. BilR and BilV have complementary regioselectivity: BilR reduces the methine bridges at C5 and C15, located between the pyrrole rings, while BilV reduces the terminal vinyl substituents at C3 and C18. Their combined action enables complete reduction to urobilinogen, and accounts for the heterogeneous urobilinoid mixture commonly observed in fecal microbial cultures. The characterization of BilV establishes that microbial bilirubin reduction is a multienzymatic process of greater complexity than previously recognized. Furthermore, the variable co-distribution of *bilR* and *bilV* across vertebrate gut microbiomes mirrors host bile pigment chemistry, suggesting co-evolution between host heme catabolism and gut microbial enzyme repertoires.

## Results

### A novel ene reductase, BilV, is genomically associated with BilR but structurally distinct

To identify additional enzymes potentially involved in bilirubin metabolism, we systematically examined the genomic neighborhoods of *bilR* across GTDB (Genome Taxonomy Database, release 226) representative genomes^12^. This analysis revealed that a subset of *bilR*-encoding species have an additional gene from a putative Old Yellow Enzyme (OYE) family in close genomic proximity to *bilR*. Based on this, we provisionally designated this gene *bilV*. BilR sequences resolve into three structurally distinct clades, BilR-short, BilR-insertion, and BilR-basal^10^, and we found that *bilV* appeared predominantly within the genomic neighborhood of BilR-insertion variants (**Fig. 1a**). A phylogenetic reconstruction of the putative BilV family showed that most but not all BilV-encoding species carry *bilR* in their genomic neighborhood (109 out of 190) (**Fig. 1b**). We observed variable gene associations including both genes being present in the same neighborhood (*Faecalibacillus intestinalis*), the genes being in separate genomic loci (*Ruminococcus gnavus* CC55_001C), and only the *bilR* gene (*Clostridium symbiosum* WAL-14163).

**Figure 1.**
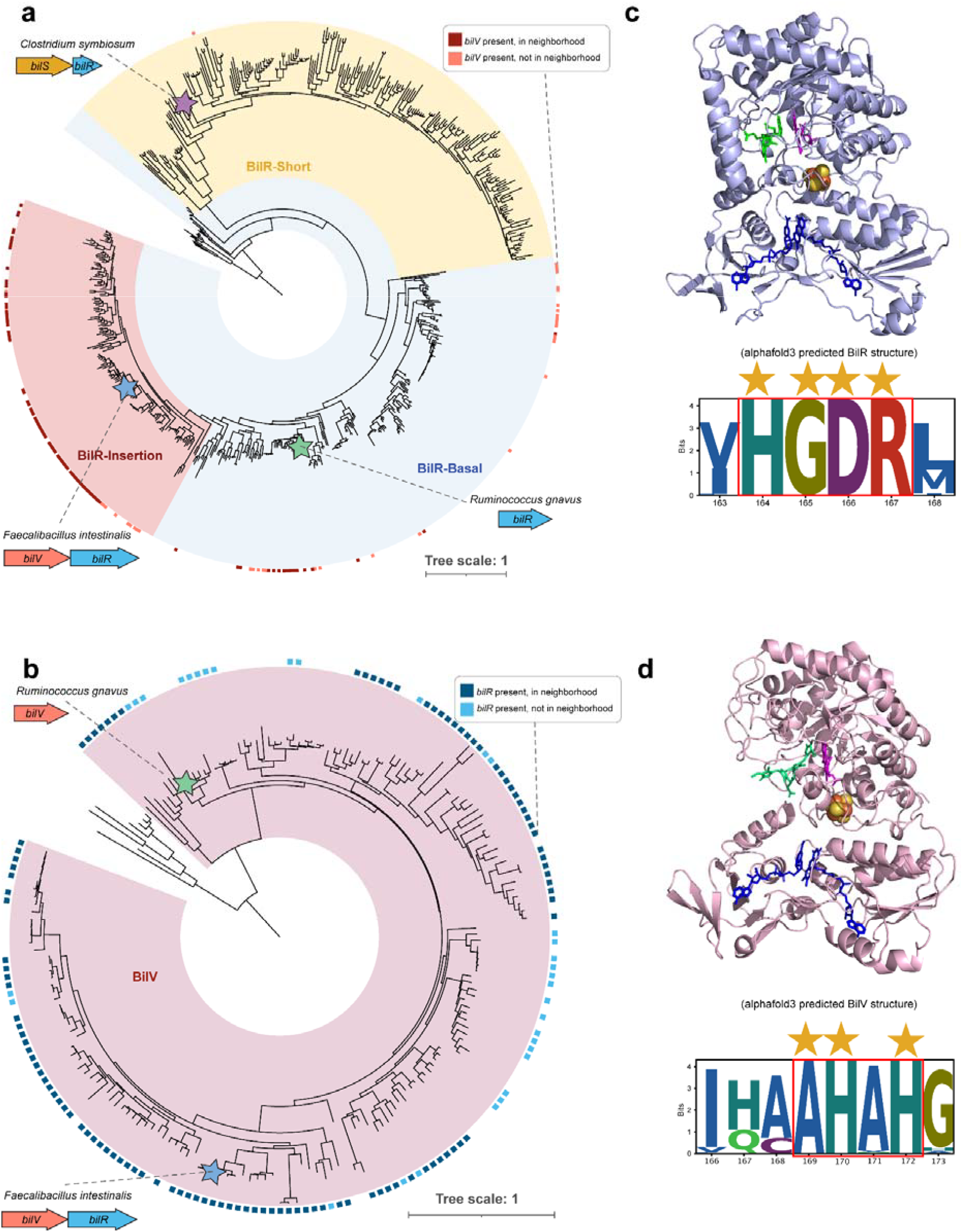
A novel ene reductase BilV is genomically associated with BilR but structurally distinct. **(a)** Maximum-likelihood phylogenetic tree of BilR protein sequences. Three BilR subtypes are highlighted: BilR-Short (yellow), BilR-Insertion (pink), and BilR-Basal (blue). Colored squares indicate species encoding both genes, with darker colors indicating presence in the same genomic neighborhood. Colored stars denote representative species functionally characterized in this study: *Clostridium symbiosum* (purple), *Ruminococcus gnavus* (green), *Faecalibacillus intestinalis* (blue). Representative gene neighborhood diagrams are shown for these species. **(b)** Maximum-likelihood phylogenetic tree of BilV protein sequences, with species marked as in (a). **(c)** AlphaFold3-predicted structure of *Ruminococcus gnavus* CC55_001C BilR showing canonical (β/α)□ TIM barrel fold. Predicted docking of bilirubin (green), and cofactors 4Fe-4S (orange and yellow), FMN (magenta) and NADH (dark blue) to BilR by AlphaFold3 are shown. The sequence logo shows the BilR canonical active site “HGDR” motif. **(d)** AlphaFold3-predicted structure of *Ruminococcus gnavus* CC55_001C BilV showing canonical (β/α)□ TIM barrel fold with predicted substrate and cofactor placements as in (c). The sequence logo of the corresponding BilV active site shows distinct conserved residues compared to BilR.

The variable gene associations prompted us to investigate whether *bilV* represents a tandem duplication of *bilR* that encodes a functionally redundant enzyme. We generated structural predictions using AlphaFold3 for both enzymes **(Fig.1c,d)**. Structural alignment of the predicted *Ruminococcus gnavus* CC55_001C BilR and BilV structures using TM-align^13^ yielded a TM-score of 0.90 and an RMSD of 2.90 Å over 648 aligned residues, indicating that BilR and BilV share the same overall fold. Despite this similarity, a comparison of the putative active site motifs in both enzymes revealed that the BilR active site contains a conserved “HGDR” motif, while the corresponding region in BilV has a distinct conserved “AHAH” motif (**Fig. 1c,d**). The structural similarity of BilR and BilV suggests that it is possible that enzymes act on similar substrates, but their divergent sequences and active site motifs suggest that they catalyze different chemical transformations.

The genomic proximity of *bilV* to *bilR* and its distinct active site architecture prompted us to hypothesize that BilV participates in bilirubin metabolism, yet it is distinct from BilR. Our initial hypothesis was that BilV might function as a urobilinogen reductase, catalyzing the further reduction of urobilinogen to stercobilinogen. However, this hypothesis was challenged by the fact that *Ruminococcus gnavus* encodes both *bilR* and *bilV*, yet is documented to produce only urobilinogen, not stercobilinogen from bilirubin in culture^3^. If BilV were a urobilinogen reductase, we would expect *R. gnavus* to convert urobilinogen further to stercobilinogen. This prompted us to consider the possibility that BilV acts on bilirubin targeting a different set of reducible bonds than BilR.

### Comparative bacterial cultivation reveals distinct bilirubin metabolites

To test whether bacterial species encoding different combinations of BilV and BilR produce distinct bilirubin-derived metabolites, we cultivated representative gut anaerobes with unconjugated bilirubin and characterized the products using complementary analytical methods. The bacteria we tested were *Clostridium symbiosum* WAL-14163 (encodes *bilR*, lacks *bilV*) and *Ruminococcus gnavus* CC55_001C (encodes both *bilR* and *bilV*). Following an 84-hour anaerobic incubation with 4.4 mg (per 100 mL of media) bilirubin, we performed the zinc acetate-urobilin fluorescence assay to detect bilirubin reduction products. Bilirubin contains four reducible carbon-carbon double bonds (**Fig. 2a**): two methine bridges connecting the pyrrole rings at C5 and C15, and two vinyl substituents on the outer rings at C3 and C18. The assay detects only molecules in which the C5 and C15 methine bridges have been reduced to single bonds, as this structural change enables zinc acetate coordination to form a fluorescent complex^3,14^. The reduction of the vinyl groups alone does not produce a signal. Both *R. gnavus* (*bilR* and *bilV*) and *C. symbiosum* (*bilR*-only) generated fluorescence significantly above the negative control (one-sided Welch’s t-test, p < 0.01; **Fig. 2b**), indicating that *bilR*-encoding species produce bilirubin-derived compounds with saturated methine bridges.

**Figure 2.**
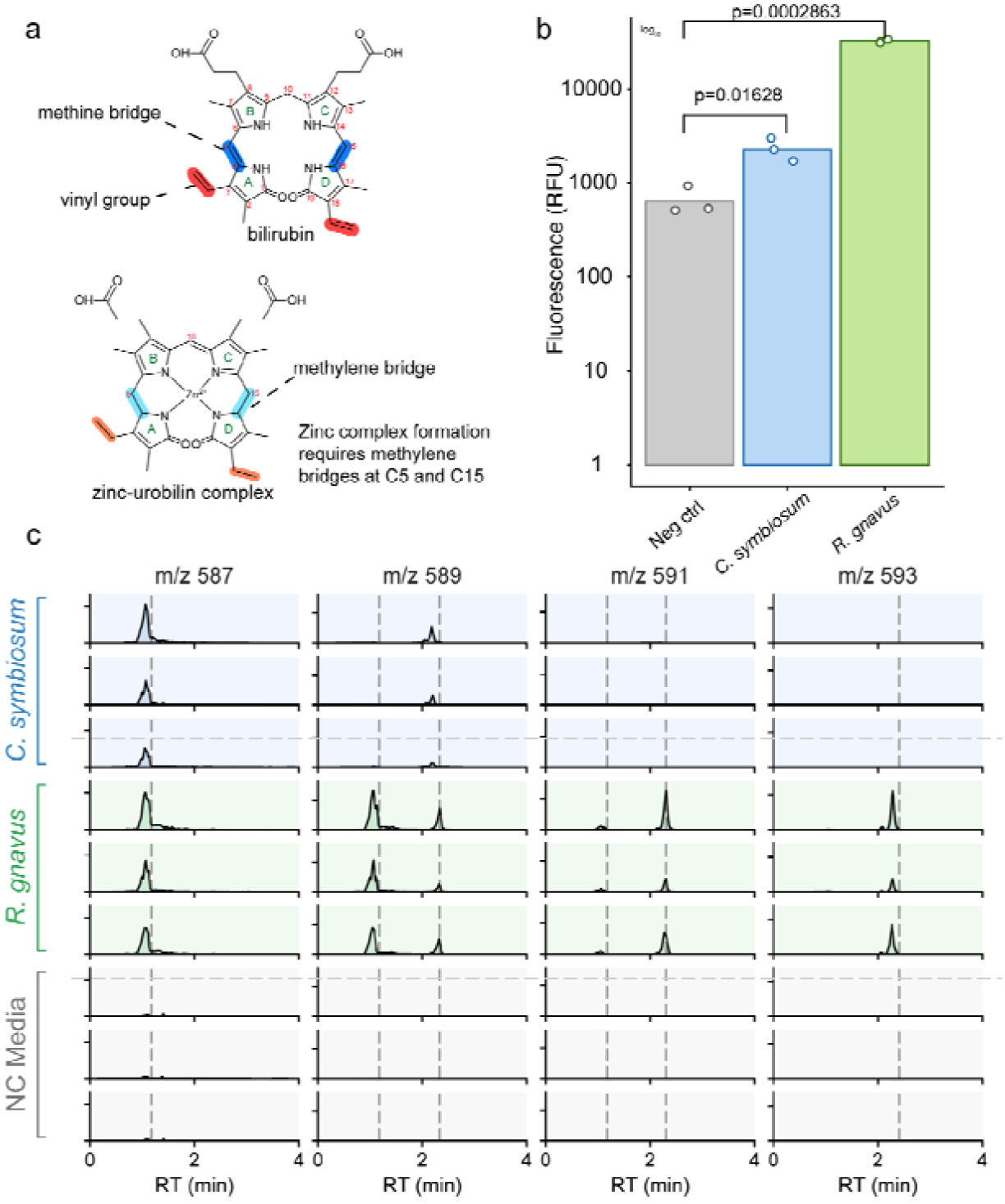
Comparative bacterial cultivation reveals distinct bilirubin metabolites. **(a)** Schematic of fluorescence assay of bilin tetrapyrroles. A Zn^2+^ ion forms a complex with the methylene bridges of the tetrapyrrole, enabling fluorescence. **(b)** zinc acetate-urobilin fluorescence assay. The bar graph shows mean fluorescence of each experimental condition (n=3 biological replicates). *C. symbiosum* (*bilR* only, blue) and *R. gnavus* (encodes both *bilV* and *bilR*, green) show strong fluorescence. Statistical comparisons between each group were made using one-tailed Welch’s t-tests. **(c)** LC-MS extracted ion chromatograms of cell cultures incubated with bilirubin.

To further characterize the bilirubin-derived products from each species, specifically reduced products, we analyzed supernatant extracts from the bacterial cultures by high-resolution LC-MS, monitoring extracted ion chromatograms (EICs) at theoretical m/z 587.2864, 589.3021, 591.3177, and 593.3334, corresponding to bilirubin reduction products with 2, 4, 6, and 8 additional hydrogens relative to bilirubin (theoretical m/z 585.2727), respectively (**Fig. 2c**). *C. symbiosum* (*bilR*-only) produced a strong signal at m/z 587.2864 (RT ~1.04 min) and a medium-intensity signal at m/z 589.3021 (RT ~2.30 min), with no detectable signal at m/z 591.3177 or 593.3334. *R. gnavus* (*bilR* and *bilV*) displayed additional products at higher m/z values, with strong signals at m/z 587.2864 (RT ~1.04 min) and m/z 589.3021 (RT ~1.11 min), a medium signal at m/z 589.3021 (RT ~2.30 min), and strong signals at m/z 591.3177 (RT ~2.30 min) and m/z 593.3334 (RT ~2.40 min), with a minor signal also detected at m/z 591.3177 (RT ~1.09 min).

These results reveal that *C. symbiosum* (*bilR*-only) and *R. gnavus* (*bilR* and *bilV*) share a prominent bilirubin-derived product at m/z 587.2864 RT ~1.04 min, while *R. gnavus* additionally produces species at higher m/z values consistent with more reduced products. Notably, the two m/z 589.3021 signals detected in *R. gnavus* at RT ~1.04 and ~2.30 min differ in retention time, indicating they are structural isomers. The structural identities of these bilirubin-derived species were determined by heterologous expression and MS/MS fragmentation analysis.

### Complementary bond specificities of BilV and BilR enable complete bilirubin reduction

To verify that the bilirubin-derived metabolites observed in bacterial cultures were specifically attributable to BilR and BilV activity, and to determine the identity of these metabolites, we heterologously expressed *R. gnavus bilR* in *E. coli*, and the *bilR-bilV* operon from *Faecalibacillus intestinalis*, incubated the cells with bilirubin or biliverdin, and characterized the reaction products by LC-MS/MS.

*E. coli* expressing *bilR* produced divinyl-urobilinogen (m/z 589.2970 [M+H]□, RT ~2.3 min), matching the m/z 589.3021 product detected in *C. symbiosum* cultures, and divinyl-urobilin (m/z 587.2809 [M+H]□, RT ~1.04 min), likely arising from spontaneous oxidation of divinyl-urobilinogen (**Fig. 3a**). MS/MS fragmentation confirmed both identities (**Fig. S1**). No signal was detected at m/z 591.3177 or 593.3334, establishing that BilR reduces the methine bridges at C5 and C15 but does not act on the vinyl groups at C3 and C18. Further LC-MS/MS analysis revealed minor products beyond divinyl-urobilinogen and divinyl-urobilin, suggesting that BilR may generate additional partially reduced intermediates from bilirubin (**Fig. S2**).

**Figure 3.**
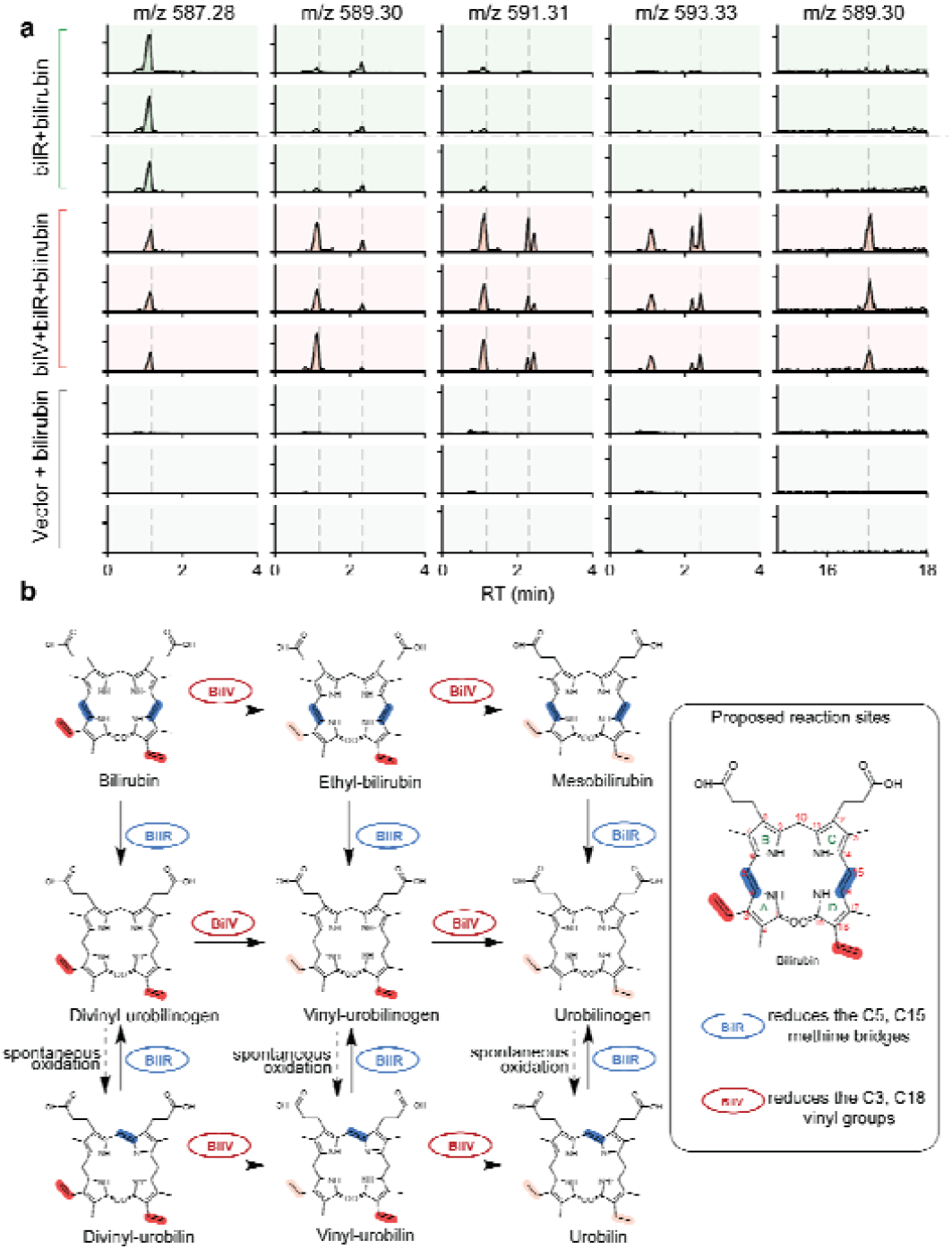
Verification of complementary BilV and BilR activity on bilirubin. **(a)** Chromatogram showing the detection of previously unidentified bilirubin metabolism intermediates at retention time 0-4 min for m/z 587.28, 589.30, 591.31, and 593.33, and then 15-18 min for 589.30, in extracts of *E. coli* expressing *bilR*-only (green), operon containing *bilV* and *bilR* (red), or vector control (gray) incubated with bilirubin. For each experimental condition, the three independent biological replicates (n□=□3) are depicted in the graph. **(b)** Proposed enzymatic pathway of bilirubin metabolism by BilV and BilR, showing complementary bond specificities for each enzyme. BilV is proposed to reduce vinyl substituents while BilR is proposed to reduce methine bridges.

*E. coli* expressing the *bilR*-*bilV* operon produced a broader product spectrum, including urobilinogen (m/z 593.3292 [M+H]□, RT ~2.43 min), urobilin (m/z 591.3145 [M+H]□, RT ~1.09 min), vinyl-urobilinogen (m/z 591.3132 [M+H]□, RT ~2.31 min), vinyl-urobilin (m/z 589.2947 [M+H]□, RT ~1.11 min), and mesobilirubin (m/z 589.3021 [M+H]□, RT ~16.8 min); empty vector controls showed no conversion (**Fig. 3a**). Compound identities were confirmed by MS/MS fragmentation (**Fig. S1**), or in the case of mesobilirubin, by retention time alone (**Fig. S3**).

When biliverdin was substituted as the substrate, *E. coli* expressing *bilR-S* from *C. difficile* alone produced divinyl-urobilinogen and divinyl-urobilin, mirroring the products observed with bilirubin, while the *bilR-bilV* operon additionally produced vinyl-reduced urobilinoids, urobilinogen, and urobilin (**Fig. S4**). These results demonstrate that BilR acts directly on biliverdin, providing a biochemical basis for its presence in avian gut microbiomes where biliverdin is the predominant bile pigment.

Because BilR alone converts bilirubin exclusively to divinyl products, the vinyl-group reduction products and mesobilirubin detected uniquely in the operon-expressing strain are attributable to BilV. Together, these results establish that BilR and BilV have complementary bond specificities, and that their combined action is necessary for complete bilirubin reduction to urobilinogen.

### Host bile pigment chemistry reflects *bilR* and *bilV* distribution across vertebrate gut microbiomes

We found *bilV* primarily within Bacillota, with the highest prevalence in the orders Erysipelotrichales (14.08%), Acholeplasmatales (2.34%), Lachnospirales (1.78%) and Oscillospirales (0.48%) (**Fig. 4a**). This distribution largely mirrored that of *bilR*, which showed highest prevalence in the same orders (Erysipelotrichales, 15.5%; Acholeplasmatales, 8.06%; Lachnospirales, 3.92%; Oscillospirales, 3.68%), though *bilR* genes were found more broadly in other orders in the Actinomycetota and Pseudomonadota phyla. *bilV* remained largely restricted to Bacillota with only minor representation in Actinomycetota (Coriobacteriales, 0.13%). The presence of *bilV* was strongly associated with *bilR* subtype (**Fig. 4b**). *bilV* co-occurred with the *bilR*-insertion variant far more frequently than with the short or basal variants. At the species level, 66.0% of *bilR*-insertion species co-encoded *bilV*, compared to 17.2% of *bilR*-basal species and only 0.4% of *bilR*-short species, with consistent proportions at the strain level (63.0%, 13.8% and 0.2%, respectively). Dufault-Thompson et al. previously showed that *bilR*-short predominates in avian gut microbiomes while *bilR*-insertion predominates in herbivores^10^. Because these two subtypes differ sharply in their genomic linkage to *bilV*, we hypothesized that *bilV* abundance varies across host species in a pattern dictated by the *bilR* subtype.

**Figure 4.**
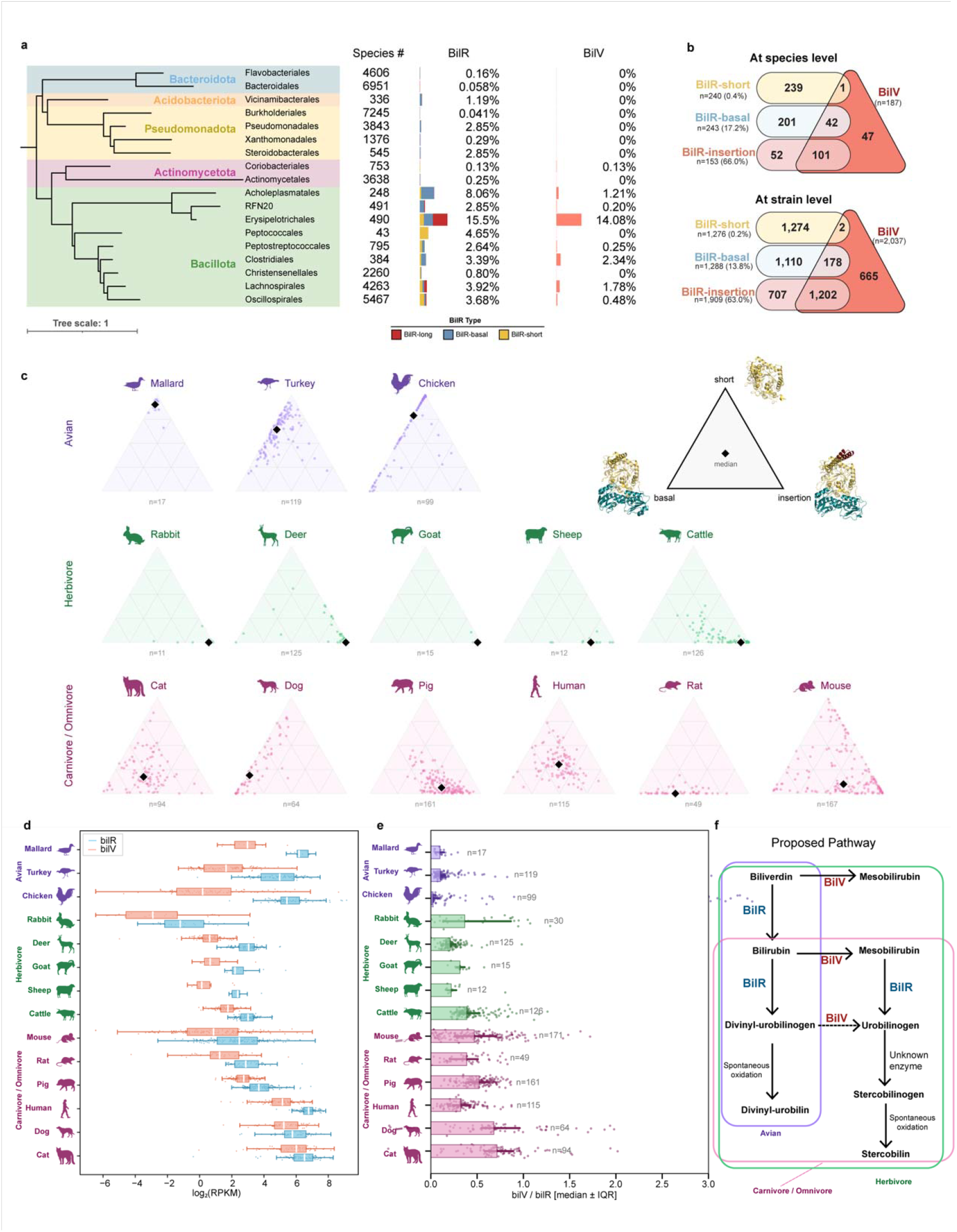
Distribution of *bilR* and *bilV* across bacterial taxa and animal gut microbiomes. **(a)** Phylogenetic distribution of *bilR* and *bilV* across bacterial orders. The cladogram shows taxonomic relationships among orders within Bacteroidota, Acidobacteriota, Pseudomonadota, Actinomycetota, Bacillota. Orders with fewer than 5 species were not displayed. 914 similar Coriobacteriales were collapsed and represented as one species. The number of species screened per order and the percentage encoding *bilR* or *bilV* are shown at right. Colored bars indicate BilR subtype (long/insertion, basal or short). **(b)** Co-occurrence of *bilV* with *bilR* subtypes at the species level (top) and strain level (bottom), shown as Venn diagrams. The numbers indicate counts of genomes encoding each *bilR* subtype alone (left), *bilV* alone (right) or both (overlap). The percentages denote the fraction of each *bilR* subtype that co-occurs with *bilV*. **(c)** Ternary plots of *bilR* subtype composition (basal, insertion, short) per sample across 14 host species grouped by dietary category: avian (purple), herbivore (green) and carnivore/omnivore (pink). Each point represents an individual metagenomic sample; black diamonds indicate per-species medians. **(d)** Paired boxplots of log_2_-transformed RPKM for *bilR* (blue) and *bilV* (red) across 14 host species. Boxes show median and interquartile range; whiskers extend to 1.5× IQR; points represent individual samples. **(e)** *bilV*/*bilR* ratio per host species. The bars indicate the median; shaded regions, the IQR; points, individual samples. A pseudocount of 0.0001 RPKM was added prior to ratio calculation to handle zero-abundance samples. **(f)** Proposed model for bilirubin reduction pathways across host species.

Vertebrates differ fundamentally in which bile pigment reaches the gut lumen, and this difference provides a physiological basis for grouping host species. Birds lack appreciable hepatic biliverdin reductase activity and excrete predominantly biliverdin, delivering little bilirubin substrate to the intestinal microbiome^11,15^. Carnivores and omnivores excrete predominantly bilirubin and produce stercobilinogen as the major fecal end product. Herbivores occupy an intermediate position, retaining low but measurable biliverdin reductase activity and excreting variable amounts of both pigments^15^. To capture this gradient in substrate availability, we grouped 14 host species into three categories: avian (chicken, turkey, mallard), herbivore (cattle, sheep, goat, deer, rabbit) and carnivore/omnivore (cat, dog, human, pig, mouse, rat). We analyzed 1,197 shotgun gut metagenomes across these 14 species (**Fig. 4c-e**). The *bilR* subtype composition agreed with previous findings: *bilR*-short accounted for 71 to 89% of total *bilR* RPKM (Reads Per Kilobase Million) in avian microbiomes, *bilR*-insertion constituted 82 to 100% in herbivores, and carnivore/omnivore species displayed more heterogeneous profiles with variable contributions from all three subtypes (**Fig. 4c**).

Consistent with the tight genomic linkage between *bilV* and *bilR*-insertion, *bilV* abundance tracked the dominant *bilR* subtype across host species (**Fig. 4d**). We detected both genes in all 1,197 samples from all 14 species. Carnivores and omnivores, which excrete bilirubin as the dominant bile pigment and harbor *bilR*-insertion at appreciable levels, carried the highest *bilV* abundance alongside high *bilR*. Herbivores carried lower but detectable *bilV* levels. Rabbits, which are obligate cecotrophs that reingest cecal pellets and effectively recycle bile pigments through a second pass of digestion and absorption, showed *bilV* near detection limits, likely because minimal bilirubin substrate reaches the distal gut microbiome. Notably, avian species carried moderately high *bilR* but markedly low *bilV*, a disparity that directly reflects the dominance of *bilR*-short, a subtype rarely associated with bilV, in these communities.

The per-sample *bilV*/*bilR* ratio captured the *bilV* deficit in avian hosts as a single metric (**Fig. 4e**). Avian species showed the lowest median ratios (0.1-fold, IQR 0-0.1, n = 231), herbivores fell in the middle (0.3-fold, IQR 0.2-0.4, n = 305) and carnivores/omnivores had the highest (0.5-fold, IQR 0.3-0.7, n = 652), indicating increasingly balanced *bilV* representation along this gradient. The balanced profile in carnivores and omnivores suggests that both the BilV-first and BilR-first pathways likely operate in these hosts, while the strongly skewed ratio in avian species points to greater reliance on the BilR-first pathway with limited vinyl-group reduction capacity. Based on these distribution patterns and the biochemical activities we established above, we propose that the relative utilization of the bilirubin reduction pathways varies across host species according to their bile pigment biochemistry and microbial *bilR* and *bilV* composition (**Fig. 4f**).

## Discussion

The characterization of BilV fills a gap in our understanding of bilirubin reduction in the gut environment. BilR directly reduces the methine bridges of bilirubin, and the co-expression data establish that BilV contributes to vinyl-group reduction of BilR-generated intermediates, providing a mechanistic basis for the heterogeneous urobilinoid mixture long observed in fecal microbial cultures. More broadly, the differential distribution of *bilV* and *bilR* subtypes across vertebrate gut microbiomes suggests that the spectrum of heme degradation end products is not fixed across vertebrate species, but shaped by the enzymatic capacity encoded in each host’s gut community.

The tight genomic linkage between *bilV* and *bilR*-insertion, but not *bilR*-short, points to a deeper functional relationship between enzyme architecture and substrate scope. *bilR*-short predominates in avian microbiomes, where host biliverdin excretion limits bilirubin availability. This subtype may therefore have experienced little selective pressure to process mesobilirubin efficiently. *bilR*-insertion, by contrast, carries an additional sequence region that may facilitate cooperation with *bilV* and dominates in hosts that excrete abundant bilirubin, environments where handling both bilirubin and the mesobilirubin intermediate would confer a clear advantage. We propose that the BilR subtypes have evolved distinct substrate preferences that mirror the bile pigment chemistry of their respective host niches, and purified-enzyme kinetic comparisons using bilirubin and mesobilirubin as substrates will test this prediction directly.

The gradient we observe, from birds (biliverdin excretors, minimal *bilV*) through herbivores (intermediate *bilV*) to carnivores and omnivores (bilirubin excretors, balanced *bilV*/*bilR*), suggests that host pigment chemistry acts as a selective filter on microbial gene content and pathway utilization. We note, however, that our model for avian hosts rests on gene-level inference rather than direct biochemical demonstration, and experiments with avian gut isolates are needed to confirm that BilR-short operates without vinyl reduction in these communities. More broadly, our survey covered only mammals and birds. Reptiles, amphibians, and fish possess distinct bile pigment profiles and gut microbial communities that could help disentangle whether *bilV* recruitment reflects substrate availability, bacterial phylogeny, or ecological niche^16,17^. Extending the comparative framework across additional vertebrate clades will clarify how host biology, bacterial adaptation, and enzyme evolution interact to shape this pathway.

The complexity of the bilirubin reduction pathway likely extends beyond what we have characterized here. Biliverdin is not fully symmetric, and the five reducible bonds: three methine bridges (C5, C10, C15) and two vinyl substituents (C3, C18), which give rise to 30 distinct intermediates between biliverdin and urobilinogen (**Figure S5**). Our data establish that BilR acts on the methine bridges and BilV reduces the vinyl groups, but the order and relative selectivity of these reductions remain unresolved. We detect vinyl-urobilin and vinyl-urobilinogen in our LC-MS profiles, yet cannot determine whether the retained vinyl group is at C3 or C18: these positional isomers are isobaric and indistinguishable without authentic standards or NMR-based structural assignment of isolated intermediates. A complete accounting of pathway intermediates will require analytical approaches capable of resolving positional isomers, alongside purified-enzyme kinetics to establish whether either enzyme exhibits a preferred bond order. The identification of BilV represents an important step toward this goal, providing the enzymatic foundation from which the full mechanistic complexity of gut microbial bilirubin catabolism can now be systematically investigated.

## Methods

### Phylogenetic analysis

BilR protein sequences were retrieved from the Genome Taxonomy Database (r226) using our previously generated profile hidden markov model (HMM) of BilR sequences^3,12^ using an e-value threshold of 1e-100. We removed 15 sequences that lacked the “HGDR” motif and 7 poorly aligned sequences. We also included 13 homologous outgroups that were not in the BilR family that we previously identified^10^. Sequences were aligned using ClustalO (v1.2.4)^18^. Maximum-likelihood phylogenetic trees were constructed using IQ-TREE (v2.3.5) with automatic model selection (ModelFinder) and 1000 ultrafast bootstrap replicates^19^. Trees were visualized and annotated using iTOL^20^.

For BilV, we constructed a profile HMM based on a curated set of sequences homologous to the *R. gnavus* BilV using hmmer^21^. We used this HMM to search for homologs in the GTDB with a cutoff of 1000 bitscore. This cutoff was empirically chosen to exclude BilR, which is homologous but functionally distinct from BilV. A phylogenetic tree was constructed using the same method as BilR.

### AlphaFold structure prediction and comparison

Protein structure predictions for BilV and BilR were generated using AlphaFold3 with default parameters^22^. Enzyme interactions with the ligands bilirubin (CID: 5280352), FMN (CID: 643976), [Fe4S4] cluster (CID: 6326971), and NADH (CID: 439153) were predicted using AlphaFold3. The BilV and BilR structures were aligned, and the TM-scores were calculated using TM-align (v20220412)^13^. The conserved BilV residues aligning to the HGDR active site motif in BilR were considered to be putative active site residues.

### Metagenomic dataset processing

Publicly available shotgun metagenomic datasets from gut samples of 14 host species were retrieved from the NCBI Sequence Read Archive (SRA), selecting species to represent three trophic groups: carnivores/omnivores (cat, dog, human, pig, mouse, rat), avian (chicken, turkey, mallard) and herbivores (cattle, sheep, goat, deer, rabbit), across 20 BioProjects (**Supplementary Table 1)**. Samples were screened for pseudoreplication arising from longitudinal sampling, multi-gastrointestinal-segment studies and crossover experimental designs. For longitudinal studies, a single time point per subject was retained: the latest time point for cattle (PRJNA1056763)^23^, human (PRJNA795985), pig (PRJEB82776) and cat (PRJEB9357)^24^, and the basal-diet period for dog (PRJNA1212753)^25^. For multi-segment studies, only caecal samples were retained from chicken (PRJNA417359)^26^, sheep (PRJNA1367213), goat (PRJNA792486)^27^ and mallard (PRJNA1269239)^28^. Young birds (day-7 and pre-weaning cohorts) were excluded from two turkey datasets (PRJNA1145617^29^, PRJNA988023)^30^. Samples with fewer than 2 million non-host read pairs were removed from one cat dataset (PRJEB79769) and one turkey dataset (PRJNA114561), and duplicate samples from repeated subjects were removed from mouse (PRJEB7759) and deer (PRJNA1083038). Technical replicates (four runs per sample) were merged for rabbit (PRJEB50625). Complete filtering criteria and per-BioProject sample counts are provided in Supplementary Table 1. After quality control, pseudoreplication filtering and depth filtering, 1,197 independent biological samples were retained across all 14 species (**Supplementary Table 2**).

Paired-end reads were quality-filtered using fastp v0.24.0 (automatic adapter detection, minimum base quality Q20, minimum read length 50 bp)^31^. Host-derived reads were removed by alignment to species-specific reference genomes using Bowtie2 v2.5.3 in sensitive mode^32^. Pre-built Bowtie2 indexes from NCBI iGenomes were used for cattle (UMD_3.1.1), pig (Sscrofa10.2), dog (Dog10K_Boxer_Tasha), mouse (GRCm38), rat (Rnor_6.0) and chicken (build3.1); the T2T-CHM13 v2.0 assembly was used for human. Custom indexes were built from NCBI RefSeq assemblies for sheep (GCF_016772045.2, ARS-UI_Ramb_v3.0), cat (GCF_018350175.1, F.catus_Fca126_mat1.0), goat (GCF_001704415.2, ARS1.2), rabbit (GCF_964237555.1, mOryCun1.1), deer (GCF_910594005.1, mCerEla1.1), mallard (GCF_047663525.1, IASCAAS_PekinDuck_T2T) and turkey (GCF_000146605.3, Turkey_5.1). Unmapped read pairs were retained for downstream quantification.

### *bilV/bilR* gene reference database and quantification

A reference database of *bilR* and *bilV* gene sequences was compiled from our curated set of homologs, comprising 4,490 *bilR* genes (1,297 basal, 1,910 insertion and 1,283 short variants) and 2,037 *bilV* genes (sequence lengths 762-3,072 bp; mean 1,789 bp). Non-host reads were aligned to this reference database using Bowtie2 v2.5.3. Per-gene read counts were extracted using samtools idxstats^33^. For samples with multiple sequencing runs, read counts were summed across technical replicates before normalisation. Gene abundance was quantified as reads per kilobase per million mapped reads (RPKM): RPKM = (read count X 10□) /(total non-host reads X gene length in bp), where total non-host reads represents the number of read pairs retained after quality filtering and host genome removal. Per-gene RPKM values were aggregated at two levels: gene class (*bilR* versus *bilV*, where *bilR* represents the sum of basal,insertion and short subtypes) and individual subtype (basal, insertion, short and *bilV*). Per-sample raw counts, SRA run accessions and RPKM values are provided in **Supplementary Table 2**.

### Anaerobic Bacterial strains and cultivation

*Clostridium symbiosum* WAL-14163 and *Ruminococcus gnavus* CC55_001C were obtained from the NIH Biodefense and Emerging Infections Research Resources Repository (BEI). These strains were inoculated from a glycerol stock and maintained under anaerobic conditions (90% N□, 5% CO□, 5% H□) in an anaerobic chamber (Coy Laboratory Products). Each strain was grown in 200 mL Brain Heart Infusion (BHI) broth (Research Products International, 32887-01-7). BHI broth was supplemented with 4.4 mg (per 100 mL of media) of bilirubin (Chem-Impex, 635-65-4) or biliverdin (Cayman Chemical Company, 856699-18-8) dissolved in dimethylsulfoxide (DMSO) and sterile filtered in the anaerobic chamber. Bacteria were incubated anaerobically at 37°C for 84 hours before the chloroform extraction and fluorescence assay were performed.

### Heterologous expression

The bilR Ruminococcus gnavus gene was PCR-amplified and cloned into pET-28a(+) (kanamycin resistance). The bilR-bilV operon from Faecalibacillus intestinalis was cloned into a pET11a vector. Both constructs were transformed into T7 Express lysY/Iq Competent E. coli (New England Biolabs, C3013I) and verified by Oxford Nanopore sequencing (Plasmidsaurus). For whole-cell biotransformation, transformed strains were inoculated from glycerol stocks into 300 mL BHI broth containing 100 µM isopropyl-B-D-1-thiogalactopyranoside (IPTG, GoldBio,367-93-1) and either 100 µg mL□^1^ carbenicillin (GoldBio, 4800-94-6) or 50 µg mL□^1^ kanamycin (Bio Basic, 70560-51-9), and shaken aerobically at 37°C for 16 hours. Cells were harvested by centrifugation (3,260 × g, 6 min) and resuspended in 200 mL BHI supplemented with bilirubin or biliverdin (4.4 mg per 100 mL, dissolved in DMSO and sterile-filtered), the appropriate antibiotic, and 100 µM IPTG. BHI medium supplemented with bilirubin or biliverdin without bacterial inoculation was included as a substrate control. Cultures were incubated anaerobically at 37°C for 24 hours, after which chloroform extraction and fluorescence assay were performed, as described below.

### Fluorescence assay

Bacterial cultures were removed from the anaerobic chamber and centrifuged at 3,260 x g for 6 minutes. The supernatant was filtered using a sterile 0.22 µm polyethersulfone (PES) filter. The filtered supernatant was extracted with 3 mL chloroform in a separatory funnel. The organic phase was collected, divided into two aliquots, and dried overnight. One aliquot was reconstituted in methanol for LC-MS/MS analysis; the other was reconstituted in 600 µL deionized water for the fluorescence assay. A 400 µL aliquot was transferred to a clean 2 mL tube to be used for the rest of the assay. 10 µL of 10% povidone-iodine solution (CVS, A47194) was added to each tube and vortexed for approximately 30 seconds. To reduce any remaining iodine, 10 µL of a 100 mM cysteine solution was added. 400 µL of Schlesinger’s reagent (545 mM zinc-acetate in methanol, ZAM) was added to each tube and vortexed. The consequent solution was used to generate 100 µL triplicates in a 96-well acrylic plate. Fluorescence was measured at 495 nm wavelength excitation and 525 nm wavelength emission at low gain using a SpectraMax M5 plate reader.

### LC-MS/MS analysis

After rehydrating the first aliquot with methanol, a 1 mL aliquot was transferred to a clean amber vial. Metabolites were analyzed by liquid chromatography-high-resolution mass spectrometry (LC-HRMS) using a Waters ACQUITY UPLC system coupled to a Bruker Daltonics maXis II quadrupole time-of-flight (Q-TOF) mass spectrometer. Samples were stored at 6°C in the autosampler prior to injection. Chromatographic separation was performed on a Phenomenex Kinetex C18 column (100 × 3.0 mm, 2.6 µm, 100 Å; core-shell) maintained at 40°C. Mobile phase A was water with 0.1% formic acid and mobile phase B was acetonitrile with 0.1% formic acid. Separation was carried out at a flow rate of 0.400 mL/min using the following gradient: 50% B (0-3 min), 50→100% B (3-18 min, linear), 100% B (18-25 min), 100→50% B (25-26 min), and re-equilibration at 50% B (26-30 min). Injection volumes were 5 µL. The mass spectrometer was operated in positive-ion electrospray ionization (ESI) mode. Source parameters: capillary voltage, 4500 V; end plate offset, −500 V; dry gas (N_2_) flow, 5.0 L/min; dry gas temperature, 220°C; nebulizer pressure, 1.0 bar. Ion transfer parameters: funnel 1 RF, 200 Vpp; funnel 2 RF, 400 Vpp; multipole RF, 200 Vpp; transfer time, 100 µs; pre-pulse storage time, 8 µs. MS1 data were acquired in profile mode over the mass range m/z 100-700. External calibration was performed using sodium formate clusters. MS/MS spectra were acquired using broadband collision-induced dissociation (BBCID).

## Data availability

The authors confirm that the data supporting the findings of this study are available within the article and its supplementary materials. All genomic data is available at the GTDB. All metagenomics data analyzed in this study are publicly available and the accession numbers can be found in **Supplementary Table 2**.

## Code availability

Relevant code for this study is available at the following GitHub repository: https://github.com/nlm-irp-jianglab/BilV-Manuscript-Bioinfo.

## Acknowledgements

This study utilized the computational resources of the NIH HPC Biowulf cluster (http://hpc.nih.gov). B.H., G.A. and M.G. are supported by R35-GM155208. This research was supported in part by the Intramural Research Program of the National Institutes of Health (NIH). The contributions of the NIH authors are considered works of the United States Government. The findings and conclusions presented in this paper are those of the authors and do not necessarily reflect the views of the NIH or the U.S. Department of Health and Human Services.We thank Sophia Levy for generating the bilR pET-28a(+) vectors.

AI-assisted tools were used for language editing and manuscript refinement. All authors reviewed and approved the final text.

## Author contributions

X.J., B.H., A.K.J. and M.G. contributed to the conceptualization of the study. M.G., X.J., and A.K.J. developed the methodology and performed validation. Formal analysis was conducted by A.K.J. and X.J., while investigation was carried out by A.K.J., G.A., M.G., A.M.C., D.L. and X.J., and data curation was performed by A.K.J. and X.J. The original draft of the manuscript was written by A.K.J., M.G., and X.J. with subsequent review and editing by K.D.T., M.G., G.A., A.K.J., X.J., and B.H. Visualization efforts were led by X.J., and A.K.J. Supervision was provided by Y.L., A.J., B.H. and X.J.

